# The effect of dream report collection and dream incorporation on memory consolidation during sleep

**DOI:** 10.1101/323667

**Authors:** Sarah F. Schoch, Maren J. Cordi, Michael Schredl, Bjöern Rasch

## Abstract

Waking up during the night to collect dream reports is a commonly used method to study dreams. This method has also been applied in studies on the relationship between dreams and memory consolidation. However, it is unclear if these awakenings influence ongoing memory consolidation processes. Furthermore, only few studies have examined if task incorporation into dreams is related to enhanced performance in the task. Here we compare memory performance in a word-picture association learning task after a night with (up to six awakenings) and without awakenings in 22 young and healthy participants. We then examine if the task is successfully incorporated into the dreams and if this incorporation is related to the task performance the next morning. We show that while the awakenings impair both subjective and objective sleep quality, these awakenings did not impair ongoing memory consolidation during sleep. When dreams were collected during the night by awakenings, memories of the learning task were successfully incorporated into dreams. No incorporation occurred in dreams collected only in the morning. Task incorporation into NREM sleep dreams, but not REM sleep dreams showed a relationship with task performance the next morning.

We conclude that the method of awakenings to collect dream reports is suitable for dream and memory studies, and is even crucial to uncover task incorporations. Furthermore, our study suggests that dreams in NREM rather than REM sleep might be related to processes of memory consolidation during sleep.

## Introduction

Current theories assume that sleep plays an active role in the process of memory consolidation. The active systems consolidation hypothesis states that memories are spontaneously reactivated and thereby redistributed between hippocampal and cortical storage sites during sleep (Born and Wilhelm, 2012). On the neural level, hippocampal reactivations occur mainly during slow-wave sleep (SWS) in rodents, and only to a lesser extent in REM sleep (Kudrimoti et al., 1999, Girardeau et al., 2017). Consequently, hippocampus-dependent declarative memories profit more from early sleep periods with high amounts of SWS (Marshall and Born, 2007). In addition, after learning a task where stimuli are linked with memory cues inducing memory reactivation by re-exposure during sleep (targeted memory reactivation) benefits are consistent when cues are presented during NREM sleep but less pronounced during REM sleep (Rasch et al., 2007, Rudoy et al., 2009, Schreiner et al., 2015).

At first glance, memory reactivations might provide an obvious link to dreaming activity (Schredl, 2017, Stickgold et al., 2001). The incorporation rate of autobiographical memories in later dreams is relatively high (Wamsley et al., 2010a, Stickgold et al., 2000, Malinowski and Horton, 2014). Dreaming occurs during both NREM and REM sleep stages, although NREM dreams are less frequent (38 - 67% vs. 75-83% in REM), shorter, less emotional and vivid (McNamara et al., 2010, Stickgold et al., 1994, Montangero, 2018). Baylor and Cavallero (2001) reported that the amount of episodic memories was higher in NREM compared to REM dream reports while there was no sleep stage dependency for semantic memories.

While waking events are clearly incorporated into dreams (Schredl and Hofmann, 2003), it is still unclear whether incorporations are related to memory consolidation. To our knowledge, only two (non-pilot) studies have examined this question using awakenings and an episodic task. In Cipolli et al. (2004), participants listened to nonsense sentences before sleep. While previously delivered sentences were incorporated more often than non-presented sentences during dream reports collected from REM sleep, incorporations made no difference in recall in the morning. In contrast, Wamsley et al. (2010b) reported that the incorporation rate during dreams in a nap positively predicts later memory performance in a spatial memory task. However, the group that dreamed about the task was very small (n = 4), and showed differences at baseline. A more general problem is that the acquisition of dreams requires repeated waking from sleep. So far it is unknown whether and how repeated collection of dream reports affects ongoing memory consolidation. Without knowing this basic effect, studies using dream collection techniques cannot be compared to most sleep and memory studies which typically examine undisturbed sleep periods.

The major aim of the current study was to examine the effect of dream report collection during sleep on memory consolidation. Additionally, we examined whether a word-image association task was incorporated into dreams and if this was related to next day memory performance. We hypothesized that repeated dream collection will disturb ongoing memory consolidation. In addition, we expected incorporations in NREM and REM sleep, but that only NREM dream incorporation will be positively related to next day memory performance.

## Methods

### Participants

Twenty-two healthy participants aged between 19 and 35 years (*M* = 23.32, SD ± 4.2) completed the whole study (12 female). They met our inclusion criteria as defined in the supplementary material and received 200 CHF as reimbursement. All participants gave written informed consent. The study was approved by the ethics committee of the Department of Psychology, University of Zurich.

### Polysomnographic set up

The polysomnographic recording consisted of electroencephalography (EEG), electrooculography (EOG), electromyography (EMG) and electrocardiography (ECG). EEG and EOG were measured with a 128 channel high density geodesic sensor net from EGI. EMG was measured with two single electrodes. ECG was measured with a singular recording from two electrodes placed on the thorax. The data went through Net Amps 300 series amplifier of EGI and was recorded and presented on the screen with the program Netstation (Version 4.5.4). Impedances were kept below 50 kΩ. Participants were woken up through an intercom system from Monacor, which allowed the experimenter to hear and talk to the participants.

### Procedures

After an adaptation night, the participants completed two experimental nights for which the participants arrived at 8 p.m. First, the polysomnography was applied. Around 9 p.m. participants started with the word-picture association learning task. After the first five blocks, they filled out two questionnaires (mood and sleep quality of the previous night), enabling a short pause between learning and recall. Then they completed the recall blocks of the task before going to bed around 11 p.m. During the experimental condition of awakenings (Session A), the participants were woken up 3 to 6 times during the night. Awakenings were based on sleep stage determined visually from the EEG. Three awakenings from NREM and three from REM sleep were prompted. Participants were immediately asked: “What went through your mind before you woke up?”. They were then asked to rate the emotionality of the content for positive and negative emotions. Participants got up at 7 a.m. in the morning and filled out a mood and sleep quality questionnaire. At the end of the session they completed the two recall blocks exactly as before sleep. In the other experimental night (nonawakening condition, B) participants were not woken up during sleep and were instructed to memorize as many dreams as possible and write them down after completing the questionnaires and memory task in the morning. Every participant remembered at least one dream. The order of awakening and non-awakening condition was counterbalanced. An overview of the procedure is depicted in Figure 1. More details on the procedure can be found in the supplementary materials.

**FIGURE 1.**
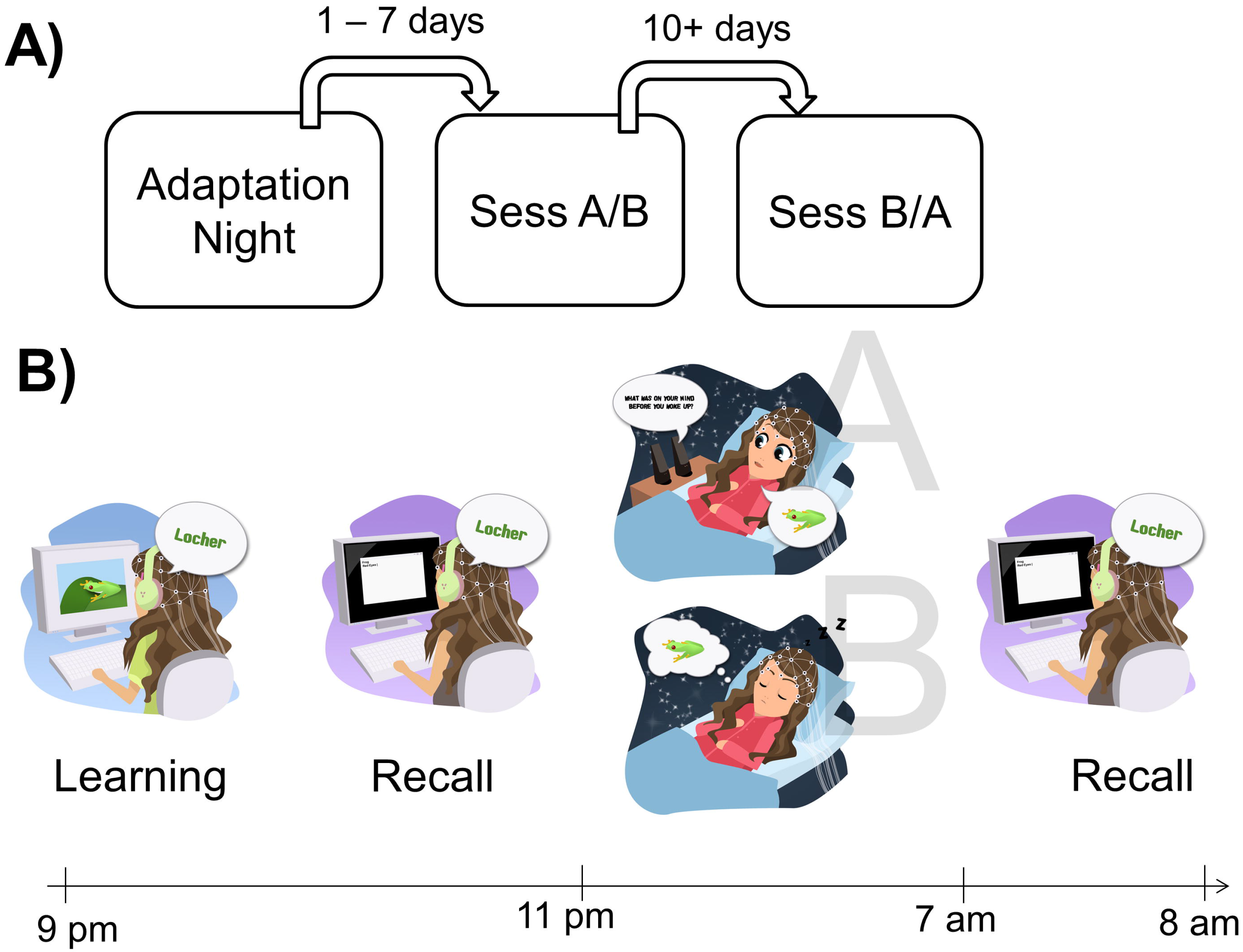

### Word-Picture Association learning Task

Memory performance was measured with a word-picture association learning task adapted from Lehmann et al. (2016b). In this task participants learned 100 neutral words, which were paired with 50 neutral and 50 positive images from 3 categories (children, sports, animals vs water, transportation and food). After rating both the words and the pictures on valence and arousal the participants tried to learn as many picture-word pairs as possible in 3 blocks. Recall happened in two blocks, once with just valence ratings and a cued recall with open answers. The percentage of the correctly remembered word-picture pairs was used as a measure for memory performance.

### Sleep and Dream Analysis

The sleep stages were scored manually using the computer software SchlafAus 1.0 (Gais, unpublished). Raters followed the rules of the Manual for the Scoring of Sleep and Associated Events from the American Academy of Sleep Medicine (Iber et al., 2007). Each dream was rated by two raters on appearance of the categories (children, sports, animals, water, transportation and food). Additionally, the dreams were rated on their realism, positive and negative feelings, number of mentioned people, acoustic perceptions, occurrences of laboratory-or experiment-related content and incorporation of the words used in the word-picture task. The interrater reliability was moderate to good (r_s_= 0.557 - 0.739, κ = 0.525 - 0.911). To operationalize to which degree the categories of the memory task were incorporated into the dreams, an incorporation score was generated for both nights and picture sets, respectively. The congruent score reflects the number of categories that had appeared in the picture set that they had seen in the task before sleep; the incongruent score reflects the number of categories of the picture set used in the other experimental night and therefore represents a baseline of the amount of task-related categories that appeared by chance in the dreams. For each reported dream the number of incorporated categories was counted (0-3) and then summed up per night. For the non-awakening condition all dreams reported in the morning were counted as one dream, for the awakening condition each report was counted as one dream. For the sleep stage-dependent analysis, only the sum of dreams that happened within the respective sleep stage was taken into account. An incorporation ratio was used to calculate the specific advantage of congruent (with the pictures presented before sleep) over incongruent items (pictures presented during the other night) during dreams in the night with awakenings and dream reports during the night (difference).

### 2.4.2 Statistical Analysis

The data was analyzed using IBM SPSS Statistics 22 (Statistical Product and Service Solutions, IBM Corp., Armonk, New York) and RStudio (R version 3.1.3; R Core Team, 2015). Statistical analysis was done with repeated measure analysis of variance (ANOVA) and one-way ANOVAs. Post-hoc analyses were corrected with Tukey’s HSD. Pairwise differences were examined using paired t-tests. For correlations, Pearson coefficients were used. Differences between the nights were examined using paired t-tests with Bonferroni corrected p-values. Significance level was set to *P* = .05.

## Results

### The effect of dream report collection on memory performance

As expected, collecting dream reports during the night strongly affected sleep quality: Compared to the night without awakenings, dream report collections during the night significantly reduced the amount of N2 and REM sleep, while it increased time spent awake after sleep onset, N1 sleep and N3 latency (see Table 1). Thus, overall sleep efficiency was significantly reduced from 94.19 ± 1.06 % in non-awakening nights to 87.34 ± 1.71 % in nights with awakenings (*P* < .001, d = 0.82, see Figure 2A).

**TABLE 1.**

|  | Awakening |  | Non-awakening |  | P | **Table 1.** |
| --- | --- | --- | --- | --- | --- | --- |
|  | M | SEM | M | SEM |  | *Comparison of* |
| Total (min) | 468.67 | ±4.74 | 454.29 | ± 8.88 | .124 | *Sleep* |
| Awake (%) | 10.43 | ± 1.53 | 2.10 | ± 0.73 | <.001* | *Characteristic* |
| N1 (%) | 7.08 | ± 0.9 | 4.61 | ± 0.7 | <.001* | *s of the* |
| N2 (%) | 52.84 | ± 1.55 | 54.72 | ± 1.22 | .27 | *Experimental* |
| N3 (%) | 13.75 | ± 0.89 | 16.93 | ± 0.64 | .007 | *Nights* |
| REM (%) | 15.9 | ± 0.88 | 21.61 | ± 1.15 | <.001* |  |
| Sleep latency (min) | 12.19 | ± 2.63 | 17.36 | ± 4.03 | .21 |  |
| SWS latency (min) | 34.43 | ± 3.88 | 16.24 | ± 1.1 | <.001* |  |
| REM latency (min) | 123.76 | ± 13.64 | 95.5 | ± 11.68 | .073 |  |
| Sleep efficiency | 87.34 | ± 1.71 | 94.19 | ± 1.06 | <.001* |  |

Note: Standard error of the means are reported. \* Significant after Bonferroni correction.

**FIGURE 2.**
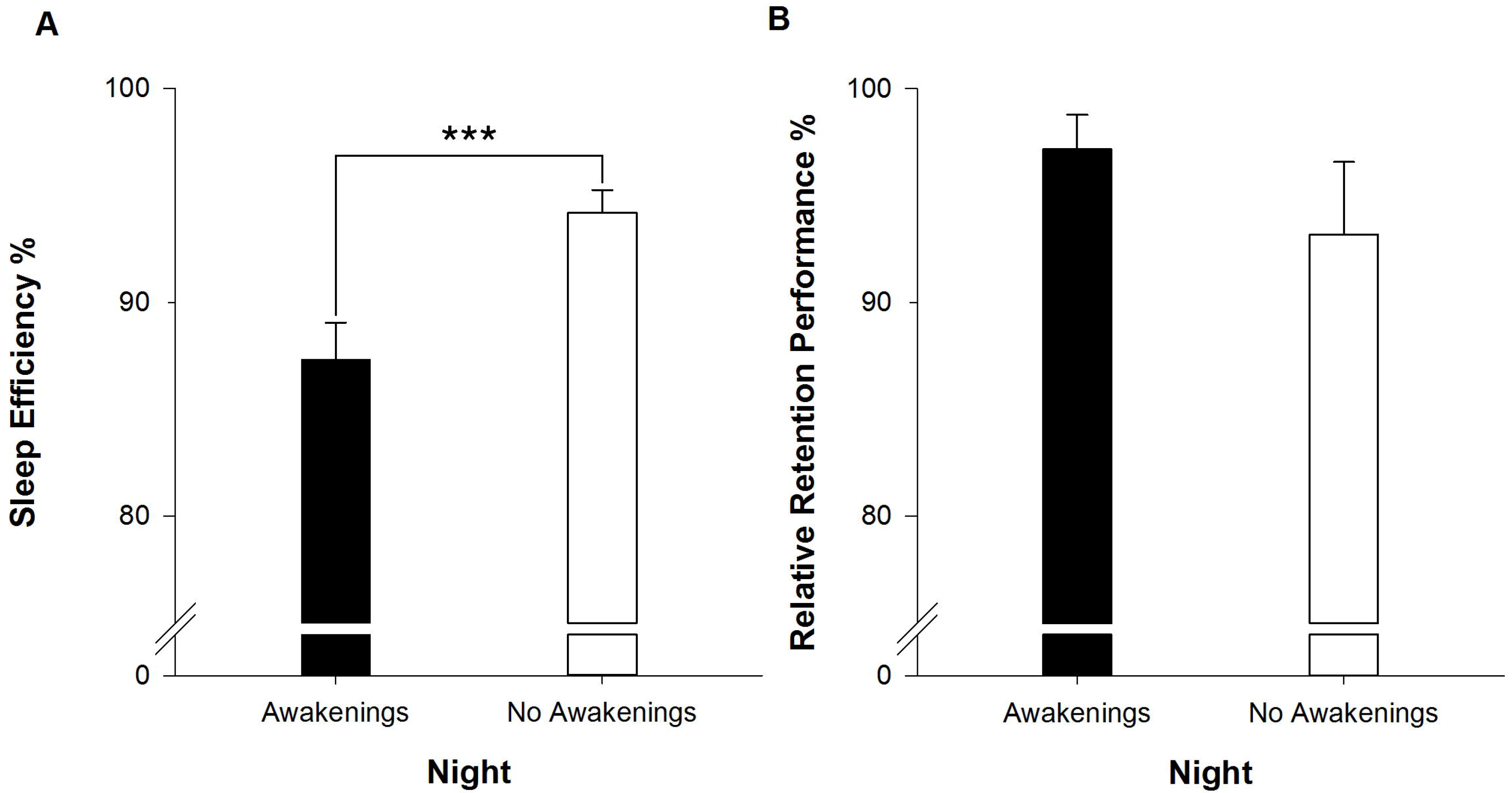

**FIGURE 3.**
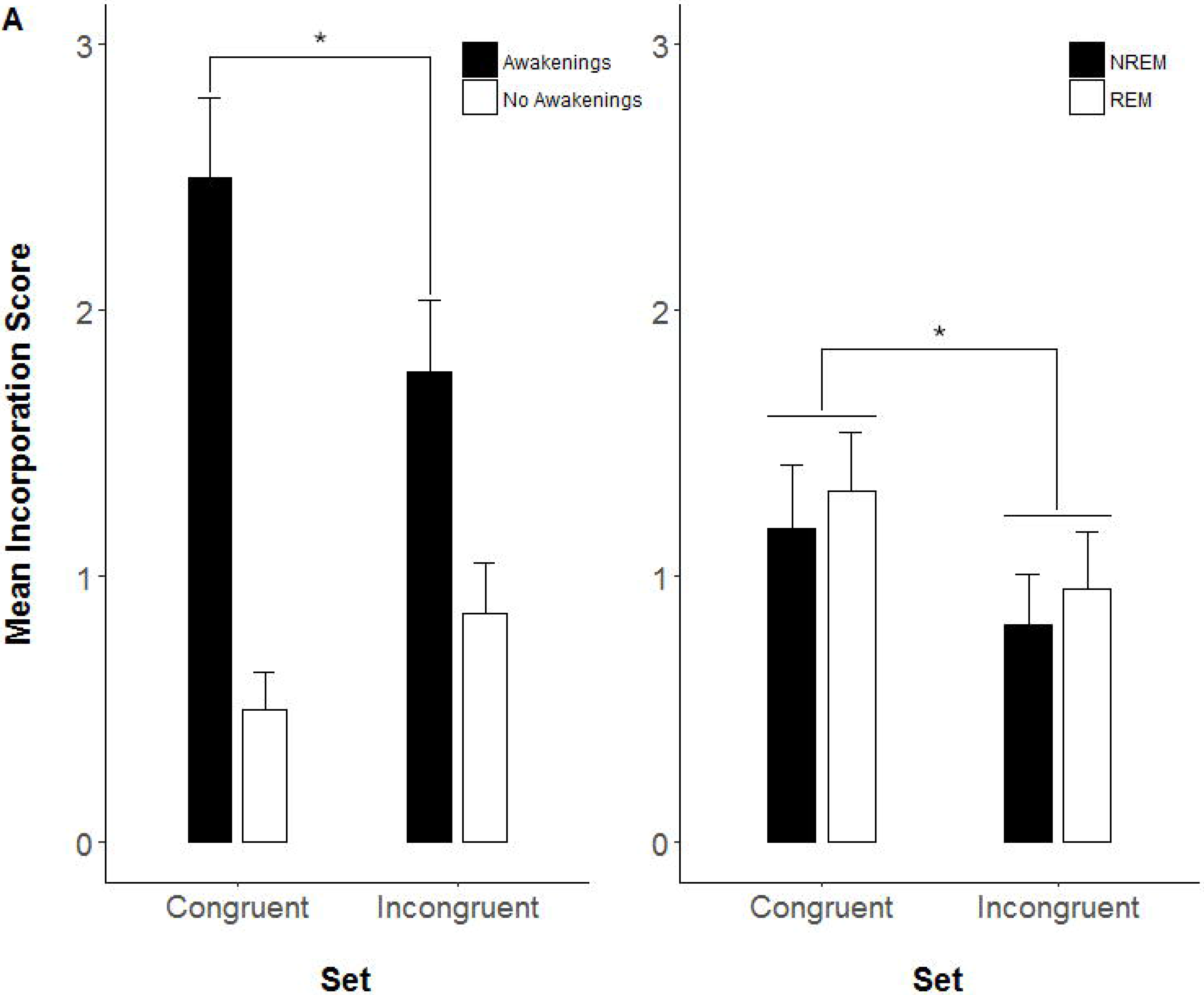

In contrast to our hypothesis and despite the strong impairment of sleep, we did not observe any significant differences between the awakening and non-awakening night on memory performance (*t*_(21)_ = 1.08, *P* = .29, d = 0.23). In the non-awakening nights, 93.19 ± 3.39 % of the images remembered in the evening were retained, with the number of images remembered in the evening before sleep set to 100%. In the night with awakenings participants remembered descriptively even more images (97.18 ± 1.61 %) (see Figure 2B). Given our sample size of n = 22, an intersession correlation of r_s_ = .27 and our alpha threshold of *P* = .05, we can exclude an effect size for independent samples with d = 0.70 or higher of the influence of awakenings on memory consolidation during sleep with a probability of 95%.

### Dream Characteristics

In the night with awakenings, participants were awakened 121 times, of which 106 lead to dream reports. 50 of these dream reports were obtained in NREM sleep (79% dream recall rate) and 56 in REM sleep (97% dream recall rate). In the night without awakenings, one morning dream diary report per participant was collected (n=22). Additional details on dream characteristics are reported in the supplementary results.

### Incorporation of task into dreams

Participants learned one image set before each experimental night (see methods and Figure 1). To test incorporation rates of images into dreams, we compared “congruent” with “incongruent” incorporations. We analyzed our data using a 2 × 2 repeated measures ANOVA with the within-subject factors night (awakening vs non-awakening) and set congruency (congruent vs incongruent). While we did not observe a main effect of set congruency (*P* > .25), we observed a significant interaction between set congruency and night (*F*_(1,21)_ = 8.1, *P* = .01, η_p_^2^ = 0.28) and a main effect of night (*F*_(1,21)_ = = 21.27, *P* < .001, η_p_^2^ = 0.5) with more incorporations in the night with awakenings. Follow-up analysis confirmed that in the night with awakenings, dream reports contained significantly more incorporations of the congruent set of categories learned before sleep (2.5 ± 0.3 incorporations) as compared to the incongruent set (1.77 ± 0.27 incorporations; *t*_(21)_ = 2.67, *P* = .014, (d = 0.57; see Figure 2A). In contrast in the nights with no awakenings, dream reports collected in the morning did not differ in the number of congruent vs. incongruent incorporations (*t*_21_ = −1.70, *P* = .10, d= 0.36, see Figure 2B.)

We further split the night with awakenings into awakenings from NREM and REM sleep stage. However, we only found a main effect of set congruency with more incorporation of the congruent set F_(1,21)_ = 7.11, *P* = .014 η_p_^2^ = 0.25, but no main effect of sleep stage or interaction between sleep stage and set congruency (all *P* ≥ .64). Thus, in both NREM and REM sleep, congruent incorporations were similarly higher as compared to incongruent incorporations.

### Relationship between Dream Incorporation and Retention Performance

We tested whether there was a positive relation between incorporation of congruent vs incongruent items of the learning task into dreams (indicated by the incorporation rate ratio between congruent and incongruent incorporations) and memory performance (measured by the relative retention score). To account for the higher chance of incorporations with more or longer dreams during REM as compared to NREM sleep, we included the amount of dreams and amount of words as covariates in a partial correlation. In accordance with our hypothesis, we observed a significant positive correlation between the ratio of congruent and incongruent dreams in NREM and overnight memory retention r =.49, *P* = .02. In contrast during REM sleep the correlation was not significant (r = -.02 *P* > .90, see Figure 4). The difference between the two correlation coefficients for NREM and REM sleep was on a trend level (z = 1.71, *P* = .087). The correlation was also not significant in the night with no awakenings, where the dreams were collected in the morning (r = .11, *P* = .61)

**FIGURE 4.**
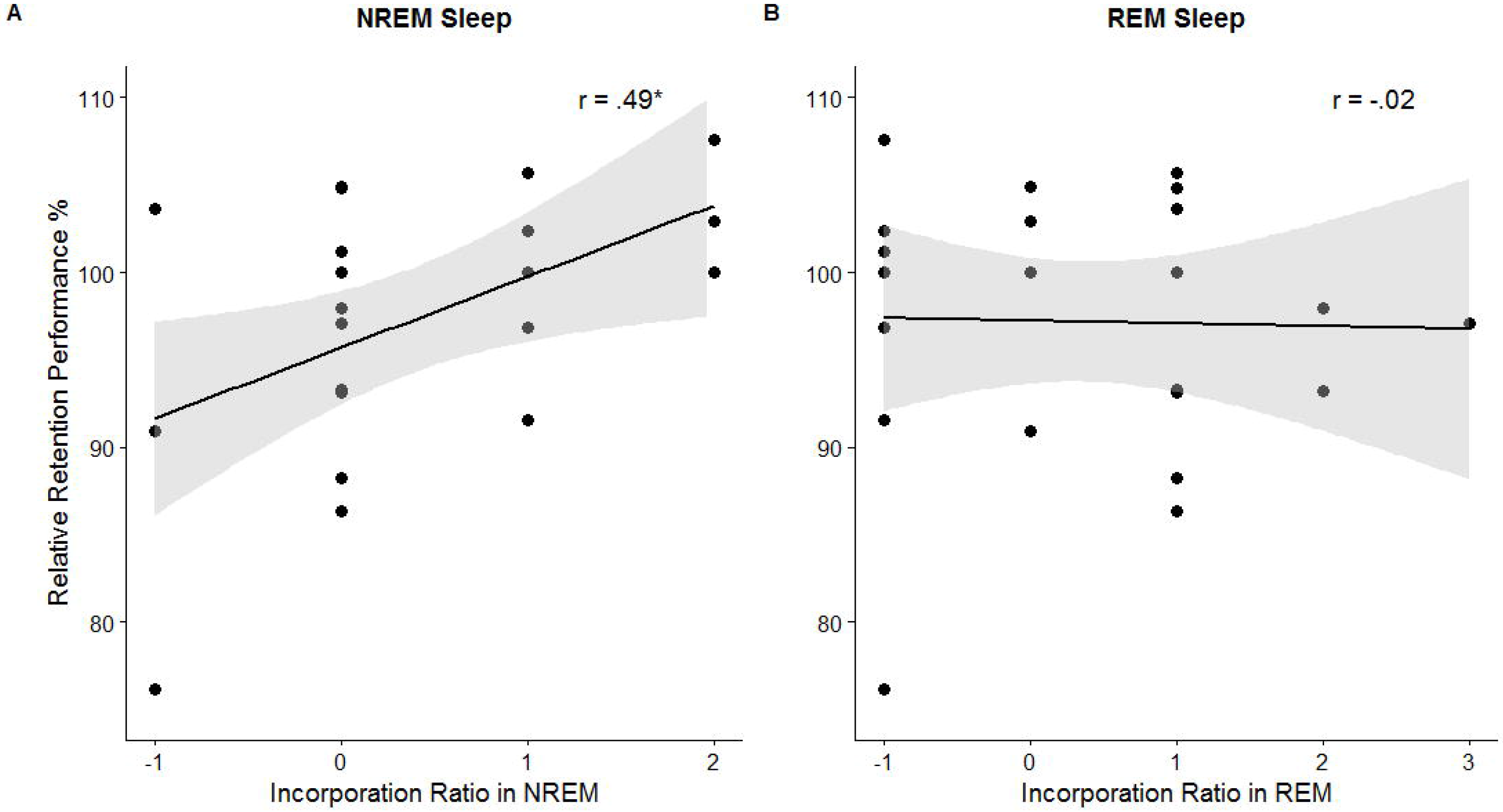

## Discussion

Our results indicate that dream report collection during sleep does not generally disturb overnight memory retention, despite impairments in sleep efficiency. Thus, memory consolidation might be comparable between nights with and without awakenings. In addition, we show that incorporation of learning stimuli into dreams reliably occurs only during dreams collected by awakenings from sleep. Finally, higher incorporation ratios of learning stimuli in NREM dreams, but not REM dreams, predicted better overnight memory retention. Our results suggest that processes of memory consolidation and reactivation during sleep might be related to dreaming during NREM sleep.

### The effect of the nocturnal awakenings on sleep-associated memory consolidation

While the awakenings impaired the objective and subjective sleep quality of the participants, they did not impair memory consolidation. Relative reductions of slow-wave sleep using procedures like the night-half paradigm (e.g. Plihal and Born, 1997) have shown that the amount of SWS might be particularly important consolidation of declarative information during sleep. Furthermore, some studies reported positive correlations between the amount of SWS and declarative memory consolidation across sleep (Backhaus et al., 2007), although this has not been consistently observed (e.g. Ackermann et al., 2015). As the amount of SWS was significantly lower in the night with awakenings, one might have expected impaired memory consolidation. However, memory performance was descriptively even better in the night with awakenings. It is possible that recalling dreams during the night represented additional processing of the task stimuli, thereby compensating for possible sleep quality impairments.

In sum, our study suggests that using up to six awakenings per night to collect dream reports does not significantly impair memory consolidation during sleep and can be used as a method to study dreams and their relationship with memory performance. However, limitations of possible additional task-related processing by repeatedly reporting dreams which might be related to the memory task apply.

### Incorporation of the task into dreams

We found that the picture set used in the task before sleep was incorporated significantly more often than the other picture set, but only if dream reports were collected by awakenings. This is an important methodological finding, as it underlines the importance of collecting dream reports via awakenings. Possibly, as dream reports in the morning only reflect a small part of the dreams that were experienced during the whole night, the remembered subset of dreams might not be representative of the whole night. A case study in one volunteer showed that both recency and intensity influenced which REM dream that was reported in the night was recalled again in the morning (Meier et al., 1968). In addition, since REM sleep stage is more prominent in the morning, it is also likely that dream reports collected in the morning rather reflect REM than NREM dreams.

While increased incorporation of the task stimuli appeared in dream reports collected via awakenings, we found no significant differences in the incorporation rate between NREM and REM dreams. According to the active system consolidation hypothesis, declarative memories are mainly re-activated during NREM sleep, while evidence for hippocampal reactivation during REM sleep is rather scarce (see Rasch and Born, 2013 for an overview). A recent study examining pattern replay in hippocampo-amygdala cell assemblies even report no signs of reactivations in REM sleep, in contrast to robust replay events during NREM sleep (Girardeau et al., 2017). Along similar lines, targeted memory reactivation during REM sleep did neither improve emotional nor neutral declarative memories, while TMR during NREM sleep improved memory for pictures (Lehmann et al., 2016a). Note that we used an almost an identical word-picture association task in the current study as Lehmann et al. (2016a). If processes of memory reactivation and dreams would be closely related, we would have expected also higher incorporation rates of our word-picture associations task in NREM. Indeed, Baylor and Cavallero (2001) reported increased incorporation of episodic memories in dream reports obtained from NREM. Possibly, the involved memory system plays a role for incorporations. Future studies will need to examine what characteristics of the task and self-relevance for the stimuli influence incorporation rates in NREM vs. REM dreams (Hoelscher et al., 1981).

### Dream incorporation and relationship with task performance

Although incorporation rates were similar between REM and NREM sleep, we found that only NREM incorporations have a positive relationship with task performance in the next morning. This is in line with the two previous studies, reporting no relation between performance and REM incorporations (Cipolli et al., 2004) and a strong association between performance and NREM incorporations (Wamsley et al., 2010b). It is possible that NREM and REM dreams reflect different mechanisms and that only NREM dreams are indicative of memory processes that take place during sleep. This also fits with the assumption of the active system consolidation hypothesis that replay mainly takes place in NREM sleep, and therefore the subjective reflection of this process also appears in this sleep stage. Our findings suggest that the association between mechanism of memory replay and dreaming might be stronger during NREM as compared to REM sleep. However, further studies are necessary to examine this notion more systematically.

### Limitations

A major limitation of our study is that we examined only dreams occurring during the first night after the memory task. According to the dream-lag effect, daily experiences get incorporated into dreams with a lag of several days (Nielsen and Powell, 1989), which might be specific to REM dreams (van Rijn et al., 2015). It is possible that incorporation into REM sleep was higher the days following the experiment, and that these incorporations would reflect ongoing memory processes. However, it is also methodologically challenging to disentangle the contributions of several nights of sleep, forgetting over time and incorporations into dreams during multiple nights to processes of memory consolidation. Another limiting factor is our sample size. While the sample size was clearly sufficient to detect differences in our within-subject design, it is not sufficient to analyze differences and associations between participants in detail (e.g. examine single items and categories in detail).

### Conclusion and future research

Here we showed that the awakenings used in dream research do not impair memory performance in an overnight task and are crucial to uncover incorporations of task into dreams. Our results support the notion that only NREM dreams might reflect ongoing memory processes, suggesting possible links between processes of memory reactivation / consolidation and dreams during NREM sleep. One might speculate that incorporation of memories during REM sleep dreams might rather support some sort of emotional processing and re-evaluation. However, the relation between processes of memory consolidation and NREM vs. REM sleep dreams clearly warrants further systematic examination.

## Acknowledgement

This work was supported by the Clinical Research Priority Program ‘Sleep and Health’ and of the University of Zurich, a grant from the Swiss National Science Foundation (SNF 100014_162388 and P0ZHP1_178697) and the European Research Council (ERC) under the European Union’s Horizon 2020 research and innovation program (grant agreement n° 677875). We thank Ilja Nefjodov and Maxime Hansen for coding the dreams and Daniel Erlacher for the support.

## References

Ackermann, S., Hartmann, F., Papassotiropoulos, A., De Quervain, D. J. F. and Rasch, B. No Associations between Interindividual Differences in Sleep Parameters and Episodic Memory Consolidation. Sleep, 2015, 38: 951–U263.

Backhaus, J., Born, J., Hoeckesfeld, R., Fokuhl, S., Hohagen, F. and Junghanns, K. Midlife decline in declarative memory consolidation is correlated with a decline in slow wave sleep. Learning & memory (Cold Spring Harbor, N.Y, 2007, 14: 336–41.

Baylor, G. W. and Cavallero, C. Memory sources associated with REM and NREM dream reports throughout the night: A new look at the data. Sleep, 2001, 24: 165–70.

Born, J. and Wilhelm, I. System consolidation of memory during sleep. Psychol Res-Psych Fo, 2012, 76: 192–203.

Cipolli, C., Fagioli, I., Mazzetti, M. and Tuozzi, G. Incorporation of presleep stimuli into dream contents: evidence for a consolidation effect on declarative knowledge during REM sleep? J Sleep Res, 2004, 13: 317–26.

Girardeau, G., Inema, I. and Buzsáki, G. Reactivations of emotional memory in the hippocampus-amygdala system during sleep. Nat Neurosci, 2017, 20: 1634.

Hoelscher, T. J., Klinger, E. and Barta, S. G. Incorporation of Concern-Related and Nonconcern-Related Verbal Stimuli into Dream Content. Journal of Abnormal Psychology, 1981, 90: 88–91.

Iber, C., Ancoli-Israel, S., Chesson, A. L. and Quan, S. F. The AASM Manual for the Scoring of Sleep and Associated Events: Rules, Terminology and Technical Specifications. In. American Academy of Sleep Medicine, Westchester, Illinois, 2007.

Kudrimoti, H. S., Barnes, C. A. and Mcnaughton, B. L. Reactivation of hippocampal cell assemblies: effects of behavioral state, experience, and EEG dynamics. J Neurosci, 1999, 19: 4090–101.

Lehmann, M., Schreiner, T., Seifritz, E. and Rasch, B. Emotional arousal modulates oscillatory correlates of targeted memory reactivation during NREM, but not REM sleep. Scientific Reports, 2016a, 6

Lehmann, M., Seifritz, E. and Rasch, B. Sleep benefits emotional and neutral associative memories equally. Somnologie, 2016b, 20: 47–53.

Malinowski, J. E. and Horton, C. L. Memory sources of dreams: the incorporation of autobiographical rather than episodic experiences. J Sleep Res, 2014, 23: 441–7.

Marshall, L. and Born, J. The contribution of sleep to hippocampus-dependent memory consolidation. Trends Cogn Sci, 2007, 11: 442–50.

Mcnamara, P., Johnson, P., Mclaren, D., Harris, E., Beauharnais, C. and Auerbach, S. Rem and Nrem Sleep Mentation. Int Rev Neurobiol, 2010, 92: 69–86.

Meier, C. A., Ruef, H., Ziegler, A. and Hall, C. S. Forgetting of Dreams in Laboratory. Percept Motor Skill, 1968, 26: 551–&.

Montangero, J. Dreaming and REM-sleep: History of a scientific denial whose disappearance entailed a reconciliation of the neuroscience and the cognitive psychological approaches to dreaming. International Journal of Dream Research, 2018, 11: 30–45.

Nielsen, T. A. and Powell, R. A. The” dream-lag” effect: A 6-day temporal delay in dream content incorporation. Psychiatric Journal of the University of Ottawa, 1989

Plihal, W. and Born, J. Effects of early and late nocturnal sleep on declarative and procedural memory. J Cognitive Neurosci, 1997, 9: 534–47.

Rasch, B. and Born, J. About Sleep’s Role in Memory. Physiol Rev, 2013, 93: 681–766.

Rasch, B., Buechel, C., Gais, S. and Born, J. Odor cues during slow-wave sleep prompt declarative memory consolidation. Science, 2007, 315: 1426–29.

Rudoy, J. D., Voss, J. L., Westerberg, C. E. and Paller, K. A. Strengthening individual memories by reactivating them during sleep. Science, 2009, 326: 1079.

Schredl, M. Is Dreaming Related to Sleep-Dependent Memory Consolidation? In), Cognitive Neuroscience of Memory Consolidation. Springer, 2017: 173–82.

Schredl, M. and Hofmann, F. Continuity between waking activities and dream activities. Conscious Cogn, 2003, 12: 298–308.

Schreiner, T., Lehmann, M. and Rasch, B. Auditory feedback blocks memory benefits of cueing during sleep. Nat Commun, 2015, 6: 8729.

Stickgold, R., Hobson, J. A., Fosse, R. and Fosse, M. Sleep, learning, and dreams: Off-line memory reprocessing. Science, 2001, 294: 1052–57.

Stickgold, R., Malia, A., Maguire, D., Roddenberry, D. and O’connor, M. Replaying the game: Hypnagogic images in normals and amnesics. Science, 2000, 290: 350–53.

Stickgold, R., Paceschott, E. and Hobson, J. A. A New Paradigm for Dream Research - Mentation Reports Following Spontaneous Arousal from Rem and Nrem Sleep Recorded in a Home Setting. Conscious Cogn, 1994, 3: 16–29.

Van Rijn, E., Eichenlaub, J.-B., Lewis, P. A. et al. The dream-lag effect: selective processing of personally significant events during rapid eye movement sleep, but not during slow wave sleep. Neurobiol Learn Mem, 2015, 122: 98–109.

Wamsley, E. J., Perry, K., Djonlagic, I., Reaven, L. B. and Stickgold, R. Cognitive Replay of Visuomotor Learning at Sleep Onset: Temporal Dynamics and Relationship to Task Performance. Sleep, 2010a, 33: 59–68.

Wamsley, E. J., Tucker, M., Payne, J. D., Benavides, J. A. and Stickgold, R. Dreaming of a Learning Task Is Associated with Enhanced Sleep-Dependent Memory Consolidation. Curr Biol, 2010b, 20: 850–55.

